# The cyclic dinucleotide 2’3’-cGAMP induces a broad anti-bacterial and anti-viral response in the sea anemone *Nematostella vectensis*

**DOI:** 10.1101/2021.05.13.443009

**Authors:** Shally R. Margolis, Peter A. Dietzen, Beth M. Hayes, Stephen C. Wilson, Brenna C. Remick, Seemay Chou, Russell E. Vance

## Abstract

In mammals, cyclic dinucleotides (CDNs) bind and activate STING to initiate an anti-viral type I interferon response. CDNs and STING originated in bacteria and are present in most animals. By contrast, interferons are believed to have emerged in vertebrates; thus, the function of CDN signaling in invertebrates is unclear. Here, we use a CDN, 2’3’-cGAMP, to activate immune responses in a model cnidarian invertebrate, the starlet sea anemone *Nematostella vectensis*. Using RNA-Seq, we found that 2’3’-cGAMP induces robust transcription of both anti-viral and anti-bacterial genes in *N. vectensis*. Many of the anti-viral genes induced by 2’3’-cGAMP are homologs of vertebrate interferon stimulated genes, implying that the interferon response predates the evolution of interferons. Knockdown experiments identified a role for NF-κB in specifically inducing anti-bacterial genes downstream of 2’3’-cGAMP. Some of these putative anti-bacterial genes were also found to be induced during *Pseudomonas aeruginosa* infection. We characterized the protein product of one of the putative anti-bacterial genes, the *N. vectensis* homolog of Dae4, and found that it has conserved anti-bacterial activity. This work suggests that a broad anti-bacterial and anti-viral transcriptional response is an evolutionarily ancestral output of 2’3’-cGAMP signaling in animals.

**Significance statement:** Cyclic dinucleotides are signaling molecules that originated in bacteria and were subsequently acquired and co-opted by animals for immune signaling. The major cyclic dinucleotide signaling pathway in mammals results in the production of anti-viral molecules called interferons. Invertebrates such as sea anemones lack interferons, and thus it was unclear whether cyclic dinucleotide signaling would play a role in immunity in these animals. Here we report that in the anemone *Nematostella vectensis*, cyclic dinucleotides activate both anti-viral and anti-bacterial immune responses, and do so through a conserved pathway. These results provide insights into the evolutionary origins of innate immunity, and suggest a broader ancestral role for cyclic dinucleotide signaling that evolved toward more specialized anti-viral functions in mammals.

## Introduction

The innate immune system is an evolutionarily ancient system that detects pathogens and initiates their elimination. In mammals, the cGAS-STING pathway is critical for sensing and responding to intracellular DNA, which is particularly important for innate responses to DNA viruses (1, 2). The sensor protein in this pathway, cyclic-GMP-AMP synthase (cGAS), is an enzyme that binds directly to cytosolic DNA and produces 2’3’-cGAMP, a cyclic dinucleotide (CDN) second messenger that binds and activates STING (3-8). Active STING uses its C-terminal tail (CTT) to recruit TBK1, which then phosphorylates and activates the transcription factor IRF3 to induce the expression of type I interferons (IFNs) (9-12). Type I IFNs are secreted cytokines that signal via JAK-STAT signaling to induce transcription of hundreds of anti-viral genes known as interferon-stimulated genes (ISGs) (13, 14). STING also activates NF-κB, MAP kinase (15), STAT6 (16), and autophagy-like pathways (17-20), as well as senescence (21) and cell death (22-26), although the mechanism of activation of these pathways, and their importance during infection, are less well understood.

Type I IFNs are thought to be a relatively recent evolutionary innovation, with identifiable interferon genes found only in vertebrates (27). In contrast, STING and cGAS are conserved in the genomes of most animals and some unicellular choanoflagellates. Remarkably, CDN to STING signaling seems to have originated in bacteria, where it may be important in bacteriophage defense (28, 29). Studies on the function of STING in animals that lack type I IFN have been mostly limited to insects, where STING seems to be protective during viral (30-33), bacterial (34), and microsporidial (35) infection. In insects, STING may promote defense through activation of autophagy (32, 35) and/or induction of NF-κB-dependent defense genes (30, 31, 34). Most ISGs are lacking from insect genomes and are not induced by CDN-STING signaling (31). In addition, the biochemical mechanisms of STING activation and signaling in insects remain poorly understood.

Biochemically, perhaps the best-characterized invertebrate STING is that of the starlet sea anemone, *Nematostella vectensis*, a member of one of the oldest animal phyla (Cnidaria). *N. vectensis* encodes a surprisingly complex genome that harbors many gene families found in vertebrates but absent in other invertebrates such as *Drosophila* (36). *N. vectensis* STING (nvSTING) and human STING adopt remarkably similar conformations when bound to 2’3’-cGAMP, and nvSTING binds to this ligand with high affinity (K_d_ < 1nM) (37). The *N. vectensis* genome also encodes a cGAS enzyme that produces 2’3’-cGAMP in mammalian cell culture (17). In vertebrates, STING requires its extended CTT to initiate transcriptional responses (38, 39); however, nvSTING lacks an extended CTT and thus its signaling mechanism and potential for inducing transcriptional responses is unclear. Based on experiments with nvSTING in mammalian cell lines, CTT-independent induction of autophagy has been proposed as the ‘ancestral’ function of STING (17), but the endogenous function of STING in *N. vectensis* has never been described.

Despite the genomic identification of many predicted innate immune genes (40, 41), few have been functionally characterized in *N. vectensis*. The sole *N. vectensis* Toll-like receptor (TLR) is reported to bind flagellin and activate NF-κB in human cell lines, and is expressed in cnidocytes, the stinging cells that define cnidarians (42). *N. vectensis* NF-κB (nvNF-κB) binds to conserved κB sites, is inhibited by *N. vectensis* IκB (43), and seems to be required for the development of cnidocytes (44). However, no activators of endogenous nvNF-κB have yet been identified. Recent work probing the putative anti-viral immune response in *N. vectensis* found that double-stranded RNA (dsRNA) injection into *N. vectensis* embryos leads to transcriptional induction of genes involved in the RNAi pathway as well as genes with homology to ISGs (45). This response is partially dependent on a RIG-I-like receptor, indicating deep conservation of anti-viral immunity (45). However, no anti-viral or anti-bacterial effectors from *N. vectensis* have been functionally tested.

Here, we characterize the response of *N. vectensis* to 2’3’-cGAMP stimulation. Similar to the response of vertebrates to 2’3’-cGAMP, we find robust transcriptional induction of putative anti-viral genes with homology to vertebrate ISGs. In addition, we observed induction of numerous anti-bacterial genes that are not induced during the vertebrate response to 2’3’-cGAMP. Although we were unable to show that the response to 2’3’-cGAMP is nvSTING dependent, we did find a selective requirement for nvNF-κB in the induction of some of the anti-bacterial genes. Many of these genes are also induced during *Pseudomonas aeruginosa* infection, suggesting a functional role in anti-bacterial immunity. We selected and characterized the anti-bacterial activity of one 2’3’-cGAMP-induced *Nematostella* gene product, domesticated amidase effector (nvDae4), a peptidoglycan cleaving enzyme that we found can kill Gram-positive bacteria. This work demonstrates an evolutionarily ancient role for 2’3’-cGAMP in the transcriptional induction of both anti-viral and anti-bacterial immunity.

## Results

### Transcriptional response to 2’3’-cGAMP in Nematostella vectensis

To assess the *in vivo* role of 2’3’-cGAMP signaling in *Nematostella vectensis*, we treated 2-week-old polyps with 2’3’-cGAMP for 24 hours and performed RNA-Seq (Fig. 1A, Fig. S1). Thousands of genes were induced by 2’3’-cGAMP, many of which are homologs of genes known to function in mammalian immunity. Despite the lack of associated gene ontology (GO) terms for many of the differentially regulated genes, unbiased GO term analysis revealed significant enrichment of immune-related terms (Fig. 1B). We also treated animals with 3’3’-linked cyclic dinucleotides, which are thought to be produced exclusively by bacteria, and which also bind to nvSTING *in vitro*, albeit at lower affinity (37). Both 3’3’-cGAMP and cyclic-di-AMP treatment also induced a smaller number of genes, although all of these genes were induced more strongly by 2’3’-cGAMP (Fig. S1). Interestingly, cyclic-di-GMP treatment led to almost no transcriptional induction, despite having relatively high affinity for nvSTING *in vitro*. This discrepancy may be due to differences in cell permeability among different CDNs, as the ligands were added extracellularly. In order to be able to perform subsequent genetic experiments, we confirmed that the immune gene induction downstream of by 2’3’-cGAMP also occurred in embryos (which are amenable to microinjection of shRNAs). Quantitative reverse transcription PCR (qRT-PCR) on 48-hour-embryos treated for 4 hours with a lower dose of 2’3’-cGAMP revealed that many immune genes were also induced at this early developmental stage after a much shorter treatment (Fig. 1C).

**Figure 1:**
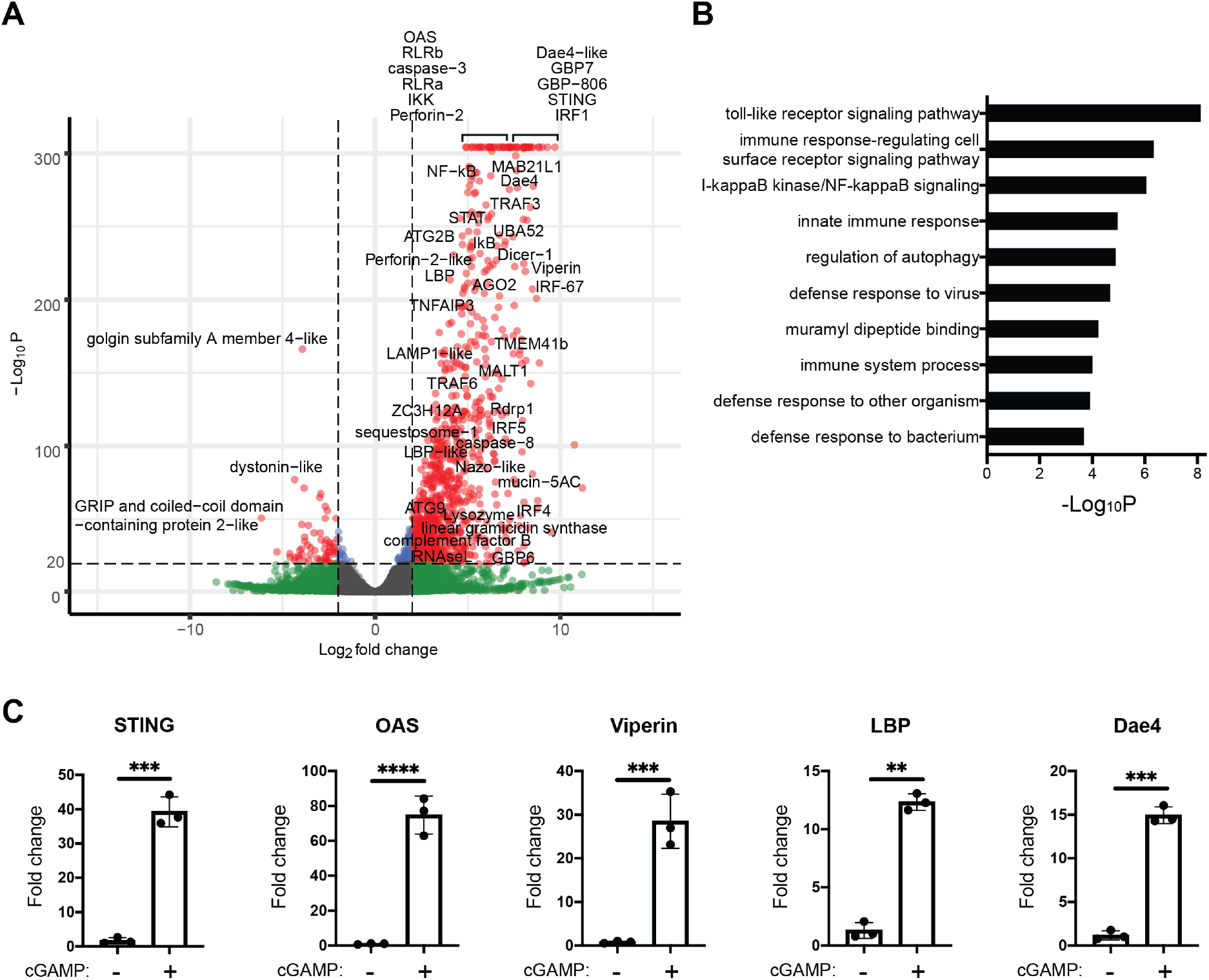
2’3’-cGAMP induces many putative immune genes in *Nematostella vectensis*. A) Volcano plot showing differential gene expression (DE) in *N. vectensis* polyps untreated vs. treated with 2’3’-cGAMP for 24 hours. A positive fold-change indicates higher expression in polyps treated with 2’3’-cGAMP. Genes of interest with homologs known to be involved in immunity in other organisms are labeled. B) Breakdown of DE genes into categories based on known GO terms. Gene set enrichment analysis shows a clear enrichment of GO terms associated with immunity. C) qRT-PCR measuring genes of interest in 48-hour-old *N. vectensis* embryos untreated or treated with 2’3’-cGAMP for 4 hours. Fold changes were calculated relative to untreated as 2^-ΔΔCt^ and each point represents one biological replicate. Unpaired t test performed on ΔΔCt before log transformation. *p ≤ 0.05; **p ≤ 0.01; ***p ≤ 0.001; ****p ≤ 0.0001.

Several interesting classes of genes were found to be upregulated in response to 2’3’-cGAMP. For example, several genes involved in the RNAi pathway were induced, including homologs of Argonaute (AGO2), Dicer, and RNA-dependent RNA polymerase (Rdrp1). In addition, many genes that are considered ISGs in mammals were also induced in *N. vectensis*, including Viperin, RNase L, 2’-5’-oligoadenylate synthase (OAS), interferon regulatory factors (IRFs), guanylate-binding proteins (GBPs), and the putative pattern recognition receptors RIG-I-like receptor a (RLRa) and RLRb. These results suggest a conserved role for 2’3’-cGAMP signaling in anti-viral immunity and ISG induction, despite an apparent lack of conservation of type I interferons in *N. vectensis*. Interestingly, we also found that many putative anti-bacterial genes were upregulated in response to 2’3’-cGAMP, including homologs of LPS-binding protein (LBP), lysozyme, perforin-2, Dae4, and mucins. These results indicate that 2’3’-cGAMP stimulation leads to a broad immune response in *N. vectensis*.

To determine whether 2’3’-cGAMP signaled via nvSTING to induce these genes, we injected shRNAs targeting nvSTING into 1-cell embryos and treated with 2’3’-cGAMP 48 hours later. We extracted RNA and performed RNA-Seq on these samples, and surprisingly, while nvSTING transcripts were reduced by ∼50%, there was no significant impact on 2’3’-cGAMP-induced gene expression (Fig. S2A, S2B). These negative results were recapitulated in numerous independent qRT-PCR and Nanostring experiments using 9 different shRNAs (3 shown in Fig. S2C). There are several possible explanations for the failure to observe a requirement for nvSTING in 2’3’-cGAMP signaling : (1) a 2-fold reduction in STING transcript levels may not result in a reduction in STING protein levels if the protein is very stable; (2) even if STING protein levels are reduced 2-fold, the reduction may not affect STING signaling due to threshold effects; or (3) nvSTING may not be required for signaling downstream of 2’3’-cGAMP due the presence of a redundant 2’3’-cGAMP sensor in *N. vectensis*. We generated an anti-nvSTING antibody to validate knockdown efficiency at the protein level, but this reagent did not appear to specifically detect nvSTING in anemone lysates. We also tested whether an nvSTING translation-blocking morpholino could inhibit induction of genes in response to 2’3’-cGAMP, but this also had no effect (Fig. S2D). Lastly, we made multiple attempts to generate nvSTING mutant animals using CRISPR, using multiple different guide RNAs, but the inefficiency of CRISPR in this organism and issues with mosaicism prevented the generation of nvSTING null animals. We previously solved the crystal structure of nvSTING bound to 2’3’-cGAMP and showed that binding occurs with high affinity (Kd < 1nM) and in a similar mode as compared to vertebrate STING (37). In addition, we found that when expressed in mammalian cells, nvSTING forms puncta only in the presence of 2’3’-cGAMP, indicating some functional change induced by this ligand (Fig. S2E). Thus, we hypothesize that 2’3’-cGAMP signals via nvSTING, but technical issues and possible redundancy with additional sensors prevent formal experimental evidence for this hypothesis.

### The N. vectensis NF-κB homolog plays a role in the 2’3’-cGAMP response

We next tested the role of conserved transcription factors that are known to function downstream of STING mammals in the *N. vectensis* response to 2’3’-cGAMP. Interestingly, many of these transcription factors are themselves transcriptionally induced by 2’3’-cGAMP in *N. vectensis* (Fig. 1A). In mammals, the transcription factors IRF3 and IRF7 induce type I IFN downstream of STING activation. While the specific function of these IRFs in interferon induction are thought to have arisen in vertebrates, other IRF family members, with conserved DNA binding residues, are present in *N. vectensis* (Fig. S3). We microinjected 1-cell embryos with short hairpin RNAs (shRNAs) targeting each of the 5 nvIRFs or a GFP control, treated with 2’3’-cGAMP, and assessed gene expression by qRT-PCR and/or Nanostring. Knockdown of IRF transcripts by 40-60% did not measurably impact gene induction by 2’3’-cGAMP (Fig. S4). We similarly tested the role of the single *N. vectensis* STAT gene, as mammalian STATs both induce anti-viral genes downstream of Type I IFN signaling, and may even be directly activated by STING (16). Similar to the nvIRFs, we did not observe a significant loss of gene induction by 2’3’-cGAMP in nvSTAT knockdown embryos by both RNA-Seq and Nanostring (Fig. S4). There are several explanations for these findings: (1) sufficient IRF or STAT protein may remain in knockdown animals to transduce the signal, either due to low efficiency of the knockdowns, or to protein stability; (2) the IRFs may act redundantly with each other, and therefore no effect will be seen in single knockdown experiments; or (3) nvIRFs and nvSTAT may not play a role in the response to 2’3’-cGAMP.

NF-κB is also known to act downstream of mammalian STING, and appears to be functionally conserved in *N. vectensis* (43). We found that NF-κB signaling components are transcriptionally induced by 2’3’-cGAMP (Fig. 1A). To test the role of nvNF-κB in the 2’3’-cGAMP response, we microinjected embryos with shRNAs targeting nvNF-κB, treated with 2’3’-cGAMP, and performed RNA-Seq (Fig. 2A). 241 genes were transcribed at significantly lower levels in the nvNF-κB knockdown embryos, and of these, 98 were genes induced by 2’3’-cGAMP. Of these genes, 40 are uncharacterized, and no GO terms were significantly enriched (data not shown). Of the induced genes that were annotated in NCBI, we noticed many were homologs of anti-bacterial proteins, including homologs of perforin-2/Mpeg-1, LPS-binding protein (LBP), linear gramicidin synthase, and mucins. We confirmed that 2’3’-cGAMP-mediated induction of these putative anti-bacterial genes was NF-κB-dependent by performing qRT-PCR and Nanostring (Fig. 2B; Fig. S5A). Of note, the induction of nvLysozyme was not nvNF-κB dependent (both by RNA-Seq and qRT-PCR; Fig. S5B), indicating either the existence of another pathway for anti-bacterial gene induction, or that our knockdown experiment was not able to affect expression of all nvNF-κB dependent genes. In addition, all of the putative anti-viral genes we examined appeared to be induced independent of nvNF-κB (Fig. S5).

**Figure 2:**
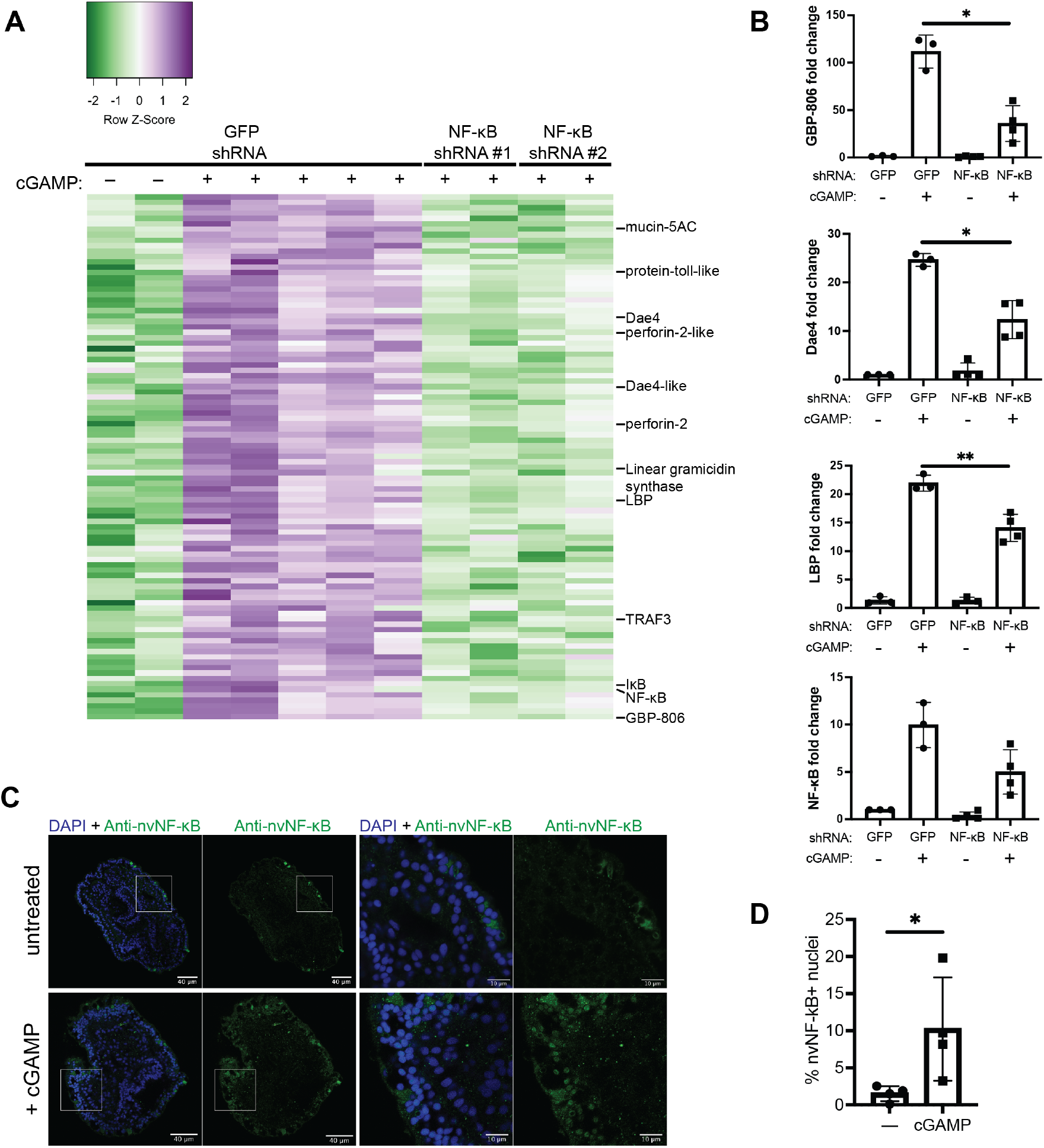
The induction of many anti-bacterial genes by 2’3’-cGAMP is nvNF-κB dependent. A) Heatmap showing all genes that are significantly (padj < 0.05, log_2_FC<-1) downregulated in 2’3’-cGAMP -treated embryos microinjected with NF-κB shRNA vs. GFP shRNA. Genes with predicted anti-bacterial function are labeled B) qRT-PCR of anti-bacterial genes in nvNF-kB shRNA or control GFP shRNA treated samples after induction by 2’3’-cGAMP. Fold change was calculated relative to untreated, GFP shRNA injected as 2^-ΔΔCt^ and each point represents one biological replicate. Unpaired t test performed on ΔΔCt before log transformation. *p ≤ 0.05; **p ≤ 0.01. C) Whole mount immunofluorescence of polyps stained with anti-nvNF-κB antiserum. Right two panels are enlargements of the boxed regions indicated in the left two panels. D) Quantification of cells with nuclear localization of nvNF-kB after treatment with cGAMP (representative images shown in C). Each point represents a single polyp, in which at least 1500 cells were analyzed. Statistical analysis was performed by unpaired t test; *p = 0.0481.

We performed BLAST searches of unannotated 2’3’-cGAMP-induced, nvNF-κB-dependent genes and identified several other genes with predicted anti-bacterial activity, including two homologs of bacterial *tae4* genes, and a putative guanylate binding protein (GBP) (*N. vectensis* LOC5515806, hereafter nvGBP-806). The Tae4 homologs had been previously identified and will be referred to as nvDae4 proteins (discussed further below; (46)). To confirm the identity of nvGBP-806 as a true GBP homolog, we performed phylogenetic analysis. We identified four conserved *N. vectensis* proteins harboring an N-terminal GBP GTPase domain with conserved GBP-specific motifs, including nvGBP-806 (Fig. S6). All of the nvGBP homologs cluster with vertebrate IFN-inducible GBPs and are themselves induced by 2’3’-cGAMP. Finally, we identified several unannotated nvNF-κB dependent, 2’3’-cGAMP-induced genes that appeared to be cnidarian-specific with no identifiable homologs in other animal phyla (Table S1).

To test directly whether nvNF-κB is activated in *N. vectensis* upon 2’3’-cGAMP treatment, we treated polyps with cGAMP and performed immunostaining for nvNF-κB (Fig. 2C). Inactive NF-κB is localized to the cytosol, and we observed sparse, cytosolic staining of ectodermal cells in untreated animals, as has been previously reported (43). In contrast, in 2’3’-cGAMP treated animals, we found many more nvNF-κB-positive cells, and in almost all of these, nvNF-κB was found in the nucleus. We performed automated quantification of nuclear nvNF-κB staining, and found that ∼3-20% of nuclei captured in our images were positive for nvNF-κB (Fig. 2D). In sum, 2’3’-cGAMP leads to nvNF-κB nuclear localization, and nvNF-κB appears to be required for expression of many putative anti-bacterial, but not anti-viral, genes. Our results demonstrate the first NF-kB agonist in *N. vectensis*, and indicate a conserved immune function for NF-kB in this organism.

### Gene induction during Pseudomonas aeruginosa challenge

In order to test whether the putative anti-bacterial, NF-κB-dependent genes are induced during bacterial infection, we infected *N. vectensis* with *Pseudomonas aeruginosa*, a pathogenic Gram-negative bacterium. *P. aeruginosa* can infect a range of hosts, including plants, mammals, and hydra (47, 48), though infections of *N. vectensis* have not previously been reported. Infection of *N. vectensis* polyps with the *P. aeruginosa* strain PA14 led to polyp death in a dose and temperature dependent manner (Fig. 3A). 48 hours after infection, we isolated RNA from infected polyps and assayed gene expression. Interestingly, nvSTING expression was induced during PA14 infection (Fig. 3B), and many of the putative anti-bacterial genes we identified as 2’3’-cGAMP-induced were also induced during infection (Fig. 3C), although this expression was not sufficient to protect from death. In addition, some putative anti-viral genes were also induced in some animals (Fig. S5B), perhaps reflecting a broader immune response in *Nematostella*. Importantly, the PA14 genome is not known to encode any proteins that produce CDNs other than c-di-GMP. Since c-di-GMP was not sufficient to robustly activate gene expression in *N. vectensis*, we believe that it is likely that the response to PA14 is independent of bacterial CDNs, although we cannot rule out an effect from PA14-produced c-di-GMP. In addition, we have no reason to believe that PA14 in activating these genes via nv-cGAS, as we do not know the activator of this enzyme. Nevertheless, taken together, these results indicate that the putative anti-bacterial genes we identified as induced by 2’3’-cGAMP are also induced after bacterial challenge.

**Figure 3:**
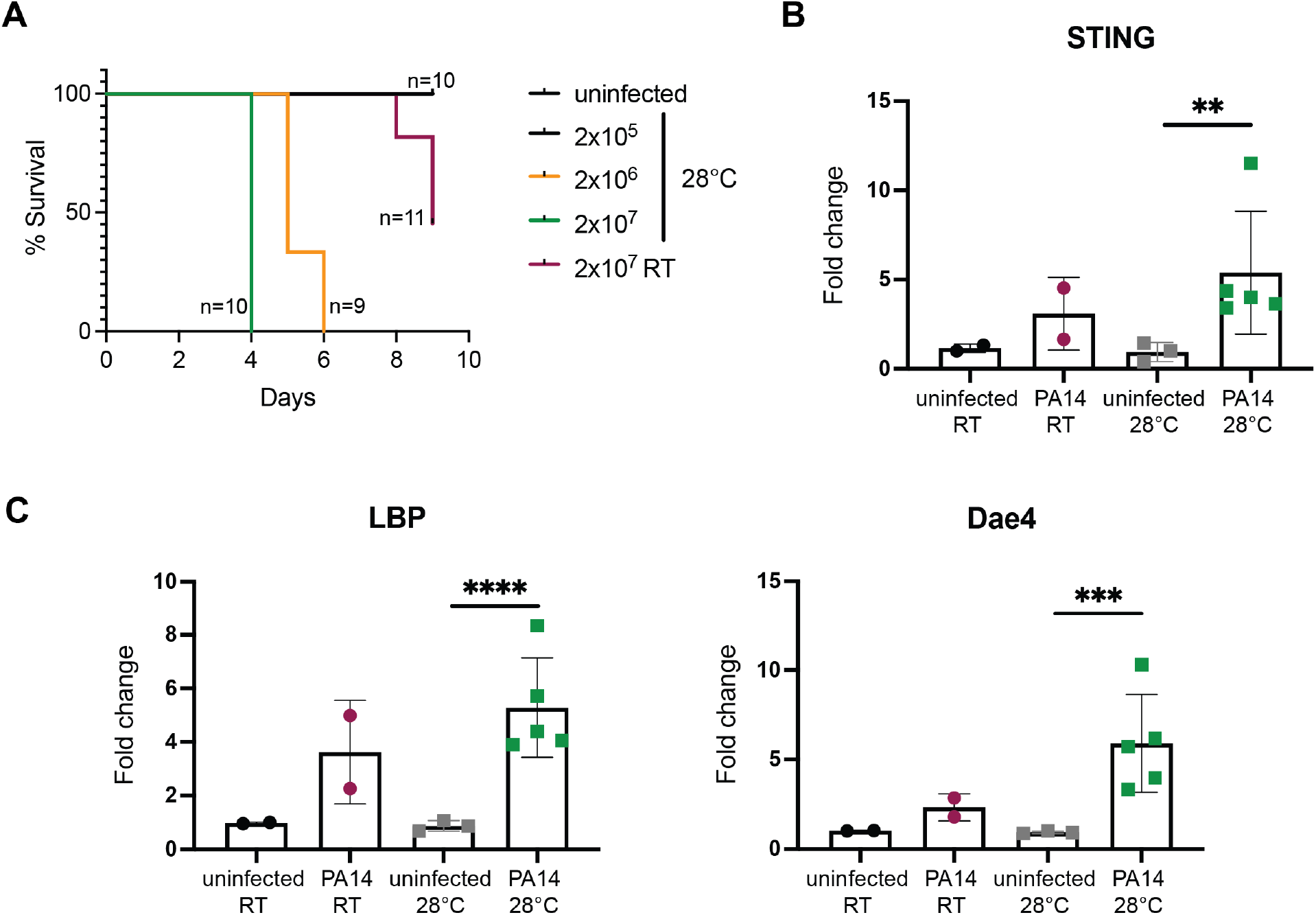
*Pseudomonas aeruginosa* infection induces putative anti-bacterial genes. A) Survival curves of *N. vectensis* polyps infected with *P. aeruginosa* at indicated dose and temperature. B+C) qRT-PCR of nvSTING (B) or putative anti-bacterial genes (C) assayed at 48 hours post *Pa* infection (2 ×10^7^ CFU/ml). Each point represents one biological replicate; unpaired t test performed on ΔΔCt before log transformation. **p ≤ 0.01; ***p ≤ 0.001; ****p ≤ 0.0001.

### nvDae4 is a peptidoglycan-cleaving enzyme with anti-bacterial activity

We decided to investigate directly whether any of the genes induced by both 2’3’-cGAMP and bacterial infection are in fact anti-bacterial. Type VI secretion amidase effector (Tae) proteins are bacterial enzymes that are injected into neighboring cells to cleave peptidoglycan, an essential component of bacterial cell walls, leading to rapid cell death (49). While the *tae* genes originated in bacteria, they have been horizontally acquired multiple times in evolution by eukaryotes, and at least one of these so-called “domesticated amidase effectors” (Daes) also has bactericidal activity (46, 50). The *N. vectensis* genome has two *tae4* homologs, both of which were upregulated by 2’3’-cGAMP in an nvNF-κB-dependent manner. However, only one of the *N. vectensis* Dae proteins is predicted to encode a conserved catalytic cysteine (46) required for peptidoglycan hydrolysis. Therefore, we focused our efforts on this homolog, which we call nvDae4 (GI: 5507694). We first tested whether nvDae4 has conserved bactericidal properties by expressing nvDae4 in *E. coli* either with or without a periplasm-targeting signal sequence and measuring bacterial growth (assessed by OD_600_) over time (Fig. 4A). *E. coli* are Gram-negative bacteria and thus have peptidoglycan compartmentalized within the periplasmic space. Consistent with the predicted peptidoglycan-cleaving function of nvDae4, only periplasmic wild-type (WT) but not catalytic mutant (C63A) nvDae4 expression led to bacterial lysis. In order to test directly whether nvDae4 cleaves peptidoglycan, we produced recombinant protein in insect cells. Since nvDae4 encodes a secretion signal, recombinant nvDae4 was secreted by the insect cells and purified from the cell supernatant. Purified nvDae4 protein was incubated with purified peptidoglycan from either *E. coli* or *Staphylococcus epidermis*. Analysis by high performance liquid chromatography (HPLC) showed that nvDae4 cleaves both Gram-negative (Fig. 4B) and Gram-positive (Fig. S7) derived peptidoglycan. Finally, we wondered whether nvDae4 could directly kill Gram-positive bacteria, as these bacteria contain a peptidoglycan cell wall that is not protected by an outer membrane and is therefore accessible to extracellular factors. We treated *B. subtilis* with recombinant nvDae4 and found that bacteria treated with WT but not C63A nvDae4 protein were killed (Fig. 4C) in a dose-dependent manner (Fig. 4D). Overall these results show that the 2’3’-cGAMP-induced protein nvDae4 is a peptidoglycan-cleaving enzyme with the capacity to kill bacteria.

**Figure 4:**
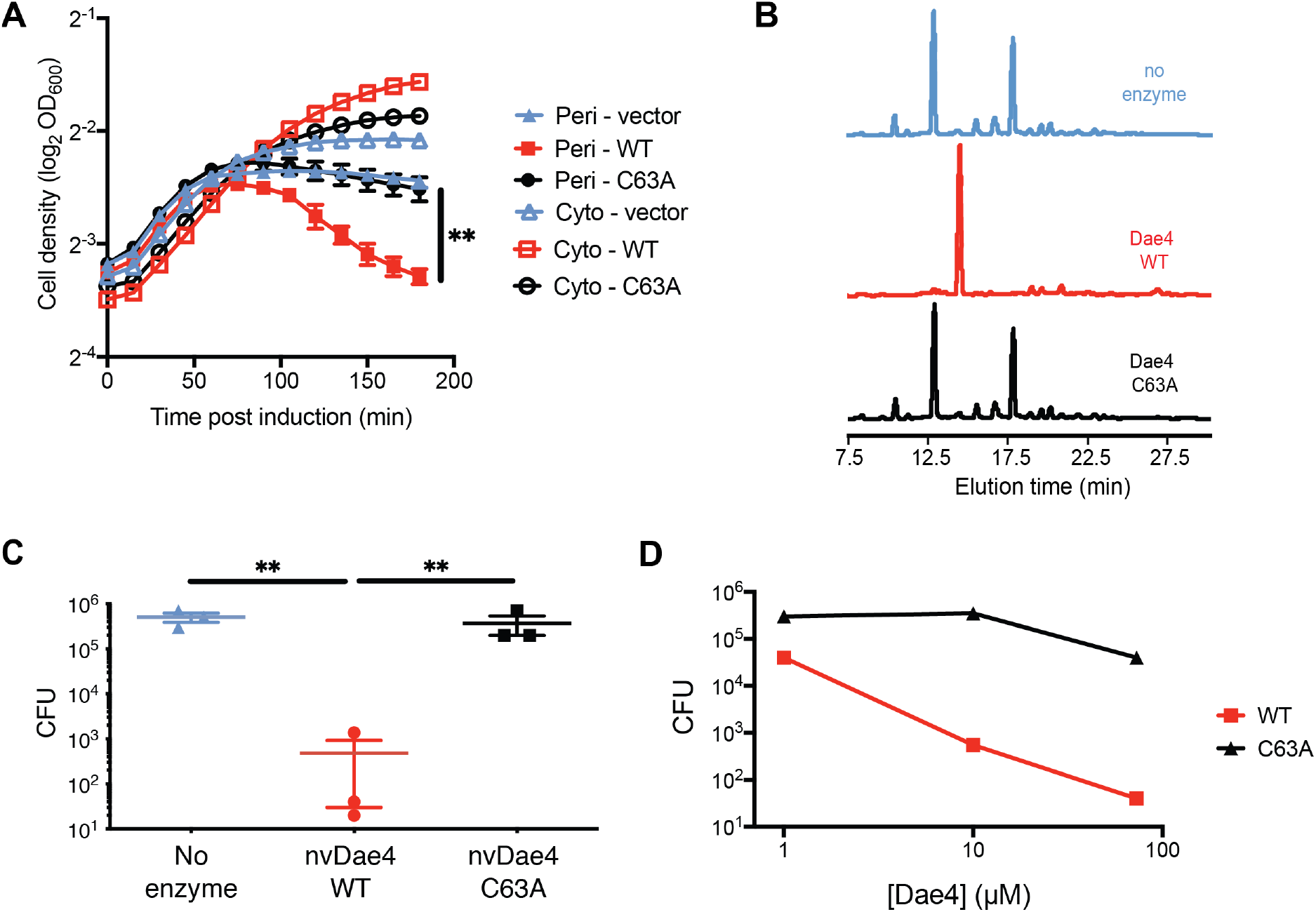
A 2’3’-cGAMP induced, nvNF-κB-dependent protein has anti-bacterial activity. A) Growth of *E. coli* expressing either periplasmic (Peri-) or cytosolic (Cyto-) nvDae4 (WT or C63A) induced with 250µM IPTG. Error bars +/-SD; n=3. Unpaired t test; **p = 0.0063. B) Partial HPLC chromatograms of *E. coli* peptidoglycan sacculi after overnight incubation with buffer only (no enzyme), or 1 µM nvDae4 WT or C63A enzyme. C) *Bacillus subtilis* CFU after 2 hour incubation with buffer alone, nvDae4 WT or catalytic mutant C63A (25 µM). Error bars +/-SEM; n=3. Unpaired t test performed on log-tranformed values; **p ≤ 0.01. D) Dose dependent killing of *B. subtilis* by WT nvDae4 enzyme (same assay as in C). Error bars +/-SD; n=2 per concentration.

## Discussion

In this study, we identified hundreds of *N. vectensis* genes that are induced by the STING ligand 2’3’-cGAMP. Despite over 600 million years of divergence and the absence of interferons, *N. vectensis* responds to 2’3’-cGAMP similarly to mammals by inducing a variety of anti-viral genes. Similarly, Lewandowska et al. (45) recently reported that *N. vectensis* responds to the synthetic double-stranded RNA poly(I:C), a viral mimic and pathogen-associated molecular pattern (PAMP). In *N. vectensis*, poly(I:C) induced both RNAi pathway components and genes traditionally thought of as vertebrate ISGs. Our combined findings indicate that the pathways linking PAMP detection to ISG expression existed prior to the vertebrate innovation of type I IFNs. Interestingly, some invertebrate species have protein-based anti-viral signaling pathways that perform similar functions to type I IFNs in vertebrates. For example, mosquito cells secrete the peptide Vago upon viral infection, which signals through the JAK-STAT pathway to activate anti-viral immunity (51). Additionally, the oyster *Crassostrea gigas* is thought to have an IFN-like system, but no secreted proteins have yet been identified in this organism (52). *N. vectensis* may also encode an undiscovered IFN-like protein; at a minimum, *N. vectensis* encodes several IRF-like genes (Fig. S3). One attractive hypothesis is that these IRFs are important for the anti-viral response of *N. vectensis*; however, we were unable to see any impact of single knockdown experiments on the induction of genes by 2’3’-cGAMP, though this may be explained by redundancy or technical limitations of our knockdown approach. Nevertheless, an important conclusion of our work is that induction of a broad transcriptional program is an ancestral function of 2’3’-cGAMP signaling, similar to what has been seen in *Drosophila* (31) and choanoflagellates (53). This ancestral transcriptional response complements an additional autophagy response to 2’3’-cGAMP that was previously reported to be induced by nvSTING in mammalian cells (17), and has now also been shown to be induced by 2’3’-cGAMP and STING in choanoflagellates (53).

We found that in addition to an anti-viral response, *N. vectensis* responds to 2’3’-cGAMP by inducing a variety of anti-bacterial genes, including lysozyme, Dae4, perforin-2-like, LPB, and GBPs. With the exception of GBPs, which have dual anti-viral and anti-bacterial activity, these anti-bacterial genes are not induced by 2’3’-cGAMP in vertebrates; thus, the anti-bacterial response appears to be a unique feature of 2’3’-cGAMP signaling in *N. vectensis*, and it will be interesting to see whether this proves true in other invertebrates, or in additional cell types or contexts in vertebrates. Indeed, a recent study found that during oral *L. monocytogenes* infection of mice, a STING-dependent and IFN-independent response helps clear bacteria. Several of the anti-bacterial genes are also induced by poly(I:C) in *Nematostella* (45), and we found that at least one putative anti-viral gene was also induced during *P. aeruginosa* infection, perhaps indicating a broader anti-pathogen response to PAMPs in *N. vectensis*. Interestingly, we found that nvNF-κB was specifically required for the induction of many of the anti-bacterial genes. This suggests that nvNF-κB activation downstream of 2’3’-cGAMP signaling may have been present in the most recent common ancestor of cnidarian and mammals, and confirms a role for nvNF-κB in *N. vectensis* immunity. Consistent with this speculation, *Drosophila* STING also appears to activate NF-κB (30, 31, 34).

To further establish that 2’3’-cGAMP induces proteins with anti-bacterial activity, we functionally characterized one 2’3’-cGAMP-induced, nvNF-κB-dependent protein, nvDae4. We found that nvDae4 is a peptidoglycan-cleaving enzyme with direct bactericidal activity against Gram-positive bacteria. Many of the 2’3’-cGAMP-induced NF-kB dependent genes are not recognizable homologs of proteins of known function; thus, they represent good candidates for the discovery of novel anti-bacterial genes in *N. vectensis*.

Using shRNAs to knockdown nvSTING failed to confirm an essential role for nvSTING in the response to 2’3’-cGAMP. However, our previous biochemical and structural studies showed nvSTING binds 2’3’-cGAMP with high affinity (K_d_ < 1nM) and in a very similar manner as vertebrate STING (37). STING is essential for the response to 2’3’-cGAMP in diverse organisms, including vertebrates, choanoflagellates (53), and insects (31). In addition, nvSTING is highly induced by 2’3’-cGAMP. So despite our negative results, we favor the idea that nvSTING is at least partially responsible for the response of *N. vectensis* to 2’3’-cGAMP. It is possible that *N. vectensis* encodes a redundant 2’3’-cGAMP sensor, but such a sensor would have had to evolve specifically in Cnidarians, or be lost independently from choanoflagellates, insects and vertebrates. It is likely that technical limitations of performing shRNA knockdowns in *N. vectensis* accounts for our inability to observe a role for nvSTING in the response to 2’3’-cGAMP, though we cannot exclude the possibility that *N. vectensis* utilizes a distinct 2’3’-cGAMP-sensing pathway.

If indeed 2’3’-cGAMP is signaling via nvSTING, this presents several mechanistic questions. First, in mammals, all known transcriptional responses downstream of STING, including those requiring NF-κB activation, require the CTT (38, 39), leading to the question of how invertebrate STING proteins, which lack a discrete CTT, can activate this pathway. Also, nvNF-κB knockdown did not impact the vast majority of 2’3’-cGAMP-induced genes, which may imply the existence of other signaling pathways downstream of nvSTING. How these unidentified pathways become activated is another interesting question and one that could also shed light on mammalian STING signaling. Finally, mammalian STING can also be activated by direct binding to bacterial 3’3’-linked CDNs (54), and nvSTING also binds to these ligands, albeit with lower affinity (37). We found that treatment of *N. vectensis* with these ligands also led to induction of many of the same genes, likely through the same pathway. This perhaps indicates a role for the nvSTING pathway in bacterial sensing, though our preliminary attempts to observe an impact of 2’3’-cGAMP-induced gene expression on bacterial colonization of *N. vectensis* were unsuccessful. Further development of a bacterial infection model for *N. vectensis* will be required to study the anti-bacterial response of this organism *in vivo*.

A crucial remaining question is what activates nvcGAS to produce 2’3’-cGAMP. Double-stranded DNA did not seem to activate this protein *in vitro* (37), but this could be due to the absence of cofactors. This protein is also constitutively active when transfected into mammalian cells, but this could be due to overexpression. Unlike human cGAS, nvcGAS does not have any clear DNA-binding domains, although this does not necessarily exclude DNA as a possible ligand. The *Vibrio* cGAS-like enzyme DncV is regulated by folate-like molecules (55), so there is a diverse range of possible nvcGAS activators. Understanding the role of CDN sensing pathways in diverse organisms can shed light on the mechanisms of evolution of viral and bacterial sensing, and on unique ways divergent organisms have evolved to respond to pathogens.

## Methods

### *Nematostella vectensis* culture and spawning

*N. vectensis* adults were a gift from Mark Q. Martindale (University of Florida) and were cultured and spawned as previously described (56). Briefly, animals were kept in 1/3x seawater (12ppt salinity) in the dark at 17°C and fed freshly hatched *Artemia* (Carolina Biological Supply Company) weekly. Spawning was induced every two weeks by placing animals at 23°C under bright light for 8 hours, followed by 2 hours in the dark, and then finally moved to the light where they were monitored for spawning. Egg masses were de-jellied in 4% L-Cysteine (pH 7-7.4) in 1/3x sea water for 10-15 minutes and washed 3 times with 1/3x sea water. Water containing sperm was added to the washed eggs and these were either used immediately for microinjection or allowed to develop at room temperature.

### CDN treatment

For the RNA sequencing experiment on polyps (Fig. 1 and Fig. S1), ∼4 week old polyps were treated in duplicate in a bath of 500μM c-di-AMP, c-di-GMP, 2’3’-cGAMP, or 3’3’-cGAMP (all InvivoGen) in 1/3x sea water for 24 hours. For remaining cGAMP treatment experiments, 50-100 48-hour old embryos were treated with 100μM 2’3’-cGAMP (InvivoGen) in 1/3x sea water for 4 hours.

### RNA sequencing

For the initial CDN treatment experiment using polyps, total RNA was extracted using Qiagen RNeasy Mini kits according to the manufacturer’s protocol. Libraries were prepared by the Functional Genomics Laboratory at UC Berkeley using WaferGen PrepX library prep kits with oligo dT beads for mRNA enrichment according to the manufacturer’s protocol, and 50 nt single-end sequencing was carried out on the HiSeq4000 (Illumina) by the Vincent J.Coates Genomics Sequencing Laboratory. For all other RNA sequencing experiments on 48 hour embryos, RNA was extracted using Trizol (Thermo Fisher Scientific) according to the manufacturer’s protocol. Libraries were prepared by the Functional Genomics Laboratory at UC Berkeley as follows: oligo dT beads from the KAPA mRNA Capture Kit (KK8581) were used for mRNA enrichment; fragmentation, adapter ligation and cDNA synthesis were performed using the KAPA RNA HyperPrep kit (KK8540). Libraries were pooled evenly by molarity and sequenced by the Vincent J.Coates Genomics Sequencing Laboratory on a NovaSeq6000 150PE S4 flowcell (Illumina), generating 25M read pairs per sample. Read quality was assessed using FastQC. Reads were mapped to the *N. vectensis* transcriptome (NCBI: GCF_000209225.1) using kallisto and differential expression was analyzed in R with DESeq2. Differential expression was deemed significant with a log_2_ fold change greater than 1 and an adjusted p-value less than 0.05. GO term analysis was performed using goseq with GO annotations from https://figshare.com/articles/dataset/Nematostella_vectensis_transcriptome_and_gene_models_v2_0/807696. The EnhancedVolcano package (https://github.com/kevinblighe/EnhancedVolcano) was used to generate volcano plots. Heatmaps are based on regularized log-transformed normalized counts and Z-scores are scaled by row. All RNA-Seq results can be found in Supplementary Dataset 1. Raw sequencing reads and normalized gene counts can be found at the NCBI GEO under accession GSE175984.

### Quantitative Real-Time PCR (qRT-PCR)

Embryos and polyps were lysed in TRIzol (Invitrogen) and RNA was extracted according to the manufacturer’s protocol. 500ng of RNA was treated with RQ1 RNase-free DNase (Promega) for and reverse transcribed with Superscript III (Invitrogen). Quantitative PCR was performed using SYBR Green (Thermo Fisher Scientific) with 0.8 μM of forward and reverse primers on a QuantStudio 5 Real-Time PCR System (Applied Biosystems) with the following cycling conditions: 50°C 2 min; 95°C 10 min; [95°C 15 sec, 60°C 1 min] x 40; 95°C 15 sec; 60°C 1 min; melt curve: step 0.075 °C/s to 95°C. Fold changes in expression levels were normalized to actin and calculated using the 2^-ΔΔCt^ method. Student’s t-tests were performed on ΔCt values. All primer sequences used in this study can be found in Supplementary Dataset 2.

### shRNA microinjection

Short hairpin RNAs for microinjection were prepared by *in vitro* transcription as previously described (57). Briefly, unique 19 nucleotide targeting motifs were identified and used to create oligonucleotides with the following sequence: T7 promoter—19nt motif—linker—antisense 19nt motif–TT. RNA secondary structure was visualized using mfold (http://www.unafold.org/mfold/applications/rna-folding-form.php) to ensure a single RNA conformation. Both sense and anti-sense oligonucleotides were synthesized and mixed to a final concentration of 25μM, heated to 98°C for 5 minutes and cooled to 24°C before use as template for *in vitro* transcription using the Ampliscribe T7-Flash Transcription Kit (Lucigen). Reactions were allowed to proceed overnight, followed by a 15 minute treatment with DNase and subsequent purification with Direct-zol™ RNA MiniPrep Plus (Zymo Research). All shRNAs used in this study can be found in Supplementary Dataset 2.

Microinjections of one-cell embryos were carried out as previously described (58). shRNAs were diluted to a concentration of 500-900 ng/μl (ideal concentrations were determined experimentally) in RNase-free water with fluorescent dextran for visualization. Injected embryos were monitored for gross normal development at room temperature and used for experiments 48 hours later unless otherwise indicated. Knockdowns for each gene were performed using at least two different shRNAs and phenotypes were confirmed in at least 3 independent experiments.

### Immunohistochemistry, imaging, and quantification

Polyps treated for 4 hours with 100μM 2’3’-cGAMP were stained for nvNF-κB as previously described (59). Briefly, polyps were fixed in 4% paraformaldehyde in 1/3x sea water overnight at 4°C with rocking, and subsequently washed 3 times with wash buffer (1× PBS, 0.2% Triton X-100). Antigen retrieval was performed by placing anemones in 95°C 5% urea for 5 minutes and allowing them to cool to room temperature before washing 3 times in wash buffer. Samples were blocked overnight at 4°C in blocking buffer (1× PBS, 5% normal goat serum, 1% bovine serum albumin, 0.2% Triton X-100). Samples were stained with anti-nvNF-κB (1:100; gift of Thomas Gilmore, Boston University) in blocking buffer for 90 minutes at room temperature and washed 4 times in wash buffer. Samples were then incubated in FITC-anti-Rabbit IgG (1:160; F9887, Sigma-Aldrich) in blocking buffer for 90 minutes at 37°C. Finally, samples were washed in wash buffer, stained with 1 μg/mL of DAPI for 10 minutes, washed again, and mounted in Vectashield HardSet Mounting medium and imaged on a Zeiss LSM 710 AxioObserver. Imaris 9.2 (Bitplane) was used to create 3D surfaces based on DAPI expression, and surface statistics were exported and analyzed in FlowJo (BD) to quantify nuclear nvNF-κB expression as previously described (60).

### Bacterial infection

Single colonies of *Pseudomonas aeruginosa* strain PA14-GFP were cultured overnight in LB with 50μg/ml of carbenicillin, centrifuged for 5 minutes at 3000 x *g*, resuspended to an OD_600_ of 0.1, 0.01, or 0.001 in 1/3x sea water and used to infect polyps, which were kept at room temperature or 28°C as indicated. Inputs were plated to calculate CFU/ml. Polyp survival was monitored daily. For expression analysis, polyps were homogenized in Trizol and RNA extraction and qPCR were performed as indicated above.

### Protein purification

nvDae4 lacking its signal peptide was cloned from cDNA. These fragments were then cloned into the pAcGP67-A baculovirus transfer vector for secreted, His-tagged protein expression. The plasmid for expressing mutant nvDae4 (C63A) was made from the pAcGP67-A-nvDae4 plasmid using Q5 site-directed mutagenesis (NEB) according to the manufactures protocol. Plasmids were transfected into Sf9 insect cells (2×10^6^ cells/ml in 2ml) using Cellfectin II Reagent (Gibco) along with BestBac 2.0 v-cath/chiA Deleted Linearized Baculovirus DNA (Expression Systems) for 6 hours, after which media was replaced and cells were left for 1 week at 25°C. Supernatants were harvested and 50 μL were used to infect 7 × 10^6^ Sf9 cells in 10ml of media for 1 week at 25°C. Supernatants containing secondary virus were harvested, tested, and used to infect High Five cells (2 L at 1.5×10^6^ cells/ml) for 72 hours at 25°C with shaking. Supernatants containing protein were harvested by centrifuging for 15 min at 600 x *g* at 4°C and subsequently passing through a 0.45 μm filter to remove all cells. Supernatants were buffered to 1x HBS (20mM HEPES pH 7.2, 150mM NaCl), mixed with 2 mL of Ni-NTA agarose were per liter, and rotated at 4°C for 2 hours. Ni-NTA resins with bound protein were collected on a column by gravity-flow and washed with 30x column volume of wash buffer (20mM HEPES, 1M NaCl, 30mM imidazole, 10% glycerol). Protein was eluted in 1 mL fractions using 1xHBS supplemented with 200mM imidazole. Buffer was exchanged to 1xHBS+ 2mM DTT using Econo-Pac10DG Desalting Prepacked Gravity Flow Columns (Bio-rad) according to the manufacturers protocol, and proteins were concentrated using 10kDa concentrators (Millipore).

### Bacterial killing assays

For expression in *E. coli*, nvDae4 WT and C63A lacking the endogenous signal sequence were cloned into the pET28a vector for inducible cytosolic expression, or the pET22b vector for inducible periplasmic expression. *E. coli* (BL21 DE3 strain) were freshly transformed with the vectors and grown overnight in LB with 50 μg/mL carbenicillin shaking at 37°C. Overnight cultures were backdiluted in LB to the same OD_600_ and grown to log phase before induction with 0.25mM IPTG. Plates were kept shaking at 37°C and OD_600_ was read every 5 minutes for 3 hours.

*Bacillus subtilis*-GFP (derivative of strain 168; BGSC accession #1A1139) was grown to log-phase in LB with 100 μg/mL spectinomycin, centrifuged, resuspended in 0.5xHBS, and incubated alone or with nvDae4 WT or C63A at indicated concentrations for 2-3 hours at 37°C. Serial dilutions were plated on LB agar with 100 μg/mL spectinomycin to determine CFU.

### Peptidoglycan cleavage assays

Peptidoglycan (PG) was purified and analyzed as previously described (50). Briefly, *Escherichia coli* BW11325 (from Carol Gross, UCSF) and *Staphylococcus epidermis* BCM060 (from Tiffany Scharschmidt, UCSF) were grown to an OD600 of 0.6, harvested by centrifugation, and boiled in SDS (4 % final concentration) for 4 hours with stirring. After washing in purified water to remove SDS, the peptidoglycan was treated with Pronase E for 2 h at 60°C (0.1 mg/ml final concentration in 10 mM Tris-HCl pH7.2 and 0.06 % NaCl; pre-activated for 2 h at 60 °C). Pronase E was heat inactivated at 100°C for 10 min and washed with sterile filtered water (5 × 20 min at 21k × *g*). PG from Gram-positive bacteria (*Se*) was also treated with 48% HF at 37°C for 48 h to remove teichoic acids, followed by washes with sterile filtered water. nvDae4 enzyme (WT and C63A) was added (1-10 µM in 10 mM Tris-HCl pH 7.2 and 0.06% NaCl) and incubated O/N at 37°C. Enzymes were heat inactivated at 100°C for 10 min. Mutanolysin (Sigma M9901, final concentration 20 µg/ml) was added to the purified peptidoglycan and incubated overnight at 37 °C. The peptidoglycan fragments were reduced, acidified, analyzed via HPLC (0.5 ml/min flow rate, 55°C with Hypersil ODS C18 HPLC column, Thermo Scientific, catalog number: 30103-254630).

## Supplemental methods

### Phylogenetic analysis

Protein sequences containing domains of interest were downloaded from NCBI, with the exception of nvGBP6 and nvGBP7, for which RNA-seq data showed additional nucleotide usage relative to the reference sequence (all sequences can be found in Supplementary Dataset 2). These were aligned using on phylogeny.fr using MUSCLE and manipulated in Geneious. For the GBP and IRF alignments, only the domain of interest was used for phylogenetic analysis. Maximum likelihood phylogenetic trees were generated with PhyML using 100 bootstrap replicates. Alignments shown were made in Geneious using Clustal Omega.

### Nanostring gene expression analysis

A custom codeset targeting 36 genes of interest and 4 housekeeping genes for normalization (Supplementary Dataset 2) was designed by NanoString Technologies (Seattle, WA) for use in nCounter XT CodeSet Gene Expression Assays run on an nCounter SPRINT Profiler (NanoString Technologies). RNA was isolated as it was for qRT-PCR experiments, and hybridized to probes according to the manufacturer’s protocol using 50ng/μL of total RNA. Quality control, data normalization, and visualization was performed in nSolver 4.0 analysis software (NanoString Technologies) according to the manufacturer’s protocol.

### Mammalian cell immunofluorescence and confocal microscopy

Glass coverslips were seeded with 293T cells and grown to ∼50% confluency. Cells were transfected for 24-48 hours with a total of 1.25 μg of DNA and 3 μL Lipofectamine 2000. Each well contained the following: pcDNA4-STING (10 ng), pEGFP-LC3 (5 ng) and either empty vector or pcDNA4 with the indicated cyclic-dinucleotide synthase (1,235 ng). Cells were washed once in PBS, fixed for 15 minutes in 4% paraformaldehyde, washed once in PBS, and permeabilized for 5 minutes in 0.5% saponin in PBS. Cells were then washed once in PBS, and treated with 0.1% sodium borohydride/0.1% saponin/PBS for 5 – 10 minutes in order to consume any remaining paraformaldehyde. Cells were then washed 3 times in PBS, and blocked with blocking buffer (1% BSA/0.1% saponin/PBS) for 45 minutes. Cells were then incubated HA antibody (1:200 dilution, Sigma 11867423001 rat IgG from Roche) in blocking buffer for one hour and washed 3 times in PBS. Each well was then incubated with 0.1% saponin/PBS and secondary antibody (1:500 dilution, Jackson ImmunoResearch, Cy3 affinipure donkey anti-rat IgG, 712-165-153) for 45 minutes. Finally, cells were washed 3 times in PBS, mounted using VectaShield with DAPI, and dried overnight. Images were acquired using a Zeiss LSM 780 NLO AxioExaminer.

## Acknowledgements

We are especially grateful to Mark Q. Martindale and Miguel Salinas-Saavedra for *N. vectensis* animals and training. We also thank Matt Gibson for sharing shRNA protocols, and Thomas Gilmore for the anti-nvNF-κB antibody. We thank members of the Vance and Barton labs for discussions, and Arielle Woznica for comments on the manuscript. This work used the Functional Genomic Laboratory and Vincent J. Coates Genomics Sequencing Laboratory at UC Berkeley, supported by NIH S10 OD018174 Instrumentation Grant. Confocal imaging experiments were conducted at the CRL Molecular Imaging Center supported by the Gordon and Betty Moore Foundation; we would like to thank Holly Aaron and Feather Ives for training and assistance. REV is an HHMI Investigator and is supported by NIH grants AI0663302, AI075039, and AI155634. SRM is supported by the National Science Foundation Graduate Research Fellowship under Grant Numbers DGE 1106400 and DGE 1752814. B.H. and S.C. were supported by funding from the NIH (R01AI132851 to S.C.), the Chan Zuckerberg Biohub, and the Sangvhi-Agarwal Innovation Award. Any opinions, findings, and conclusions or recommendations expressed in this material are those of the author(s) and do not necessarily reflect the views of the National Science Foundation, HHMI, or the National Institutes of Health.

## Figure Legends

**Figure S1:**
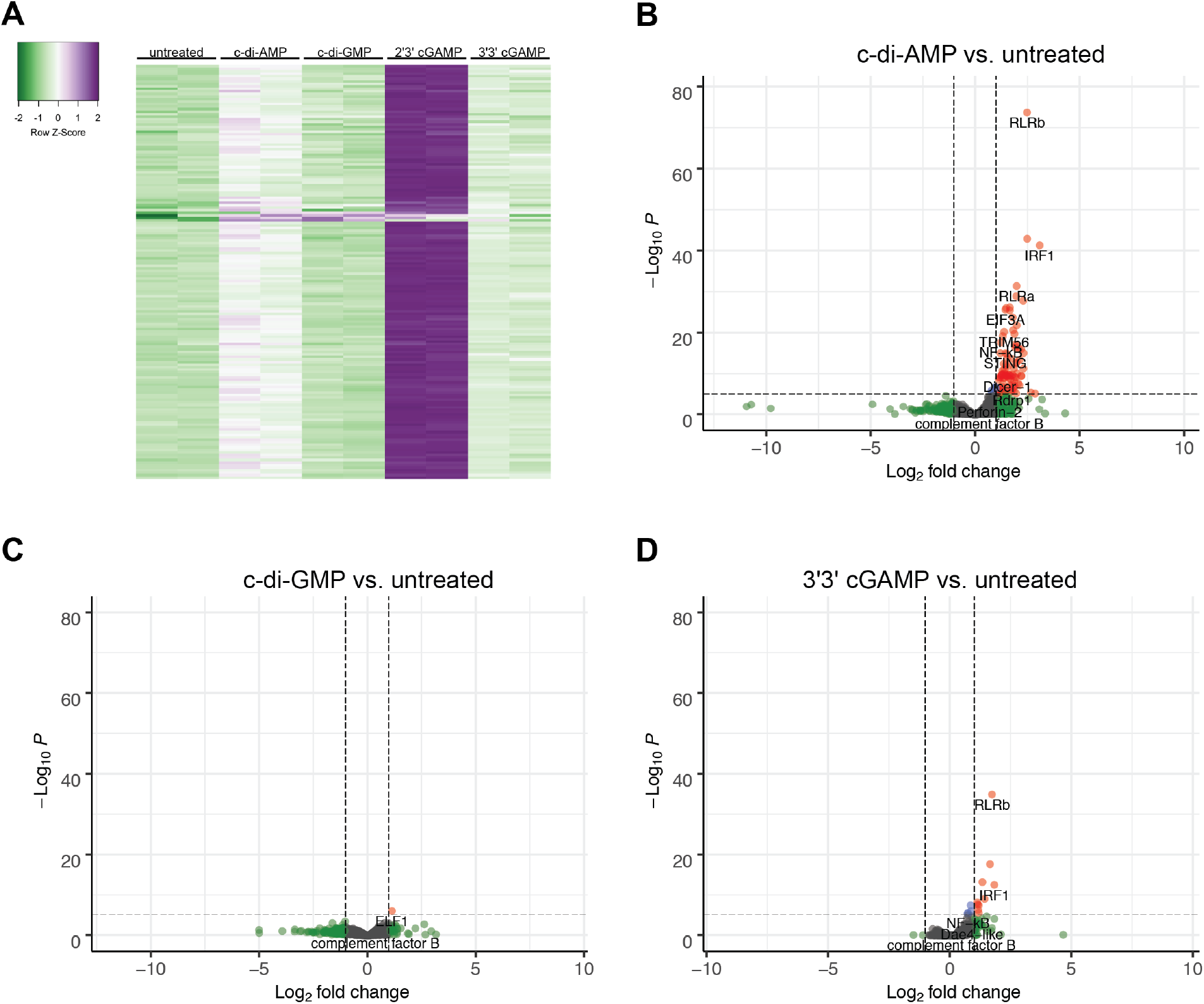
Treatment with other CDNs leads to some gene induction. A) Heatmap showing differentially expressed genes in response to c-di-AMP, c-di-GMP, and 3’3’-cGAMP. Almost all of these are also significantly induced by 2’3’-cGAMP. B-D) Volcano plots of differential gene expression in *N. vectensis* polyps untreated vs. treated with cyclic-di-AMP (B), cyclic-di-GMP (C) and 3’3’-cGAMP (D) for 24 hours.

**Figure S2:**
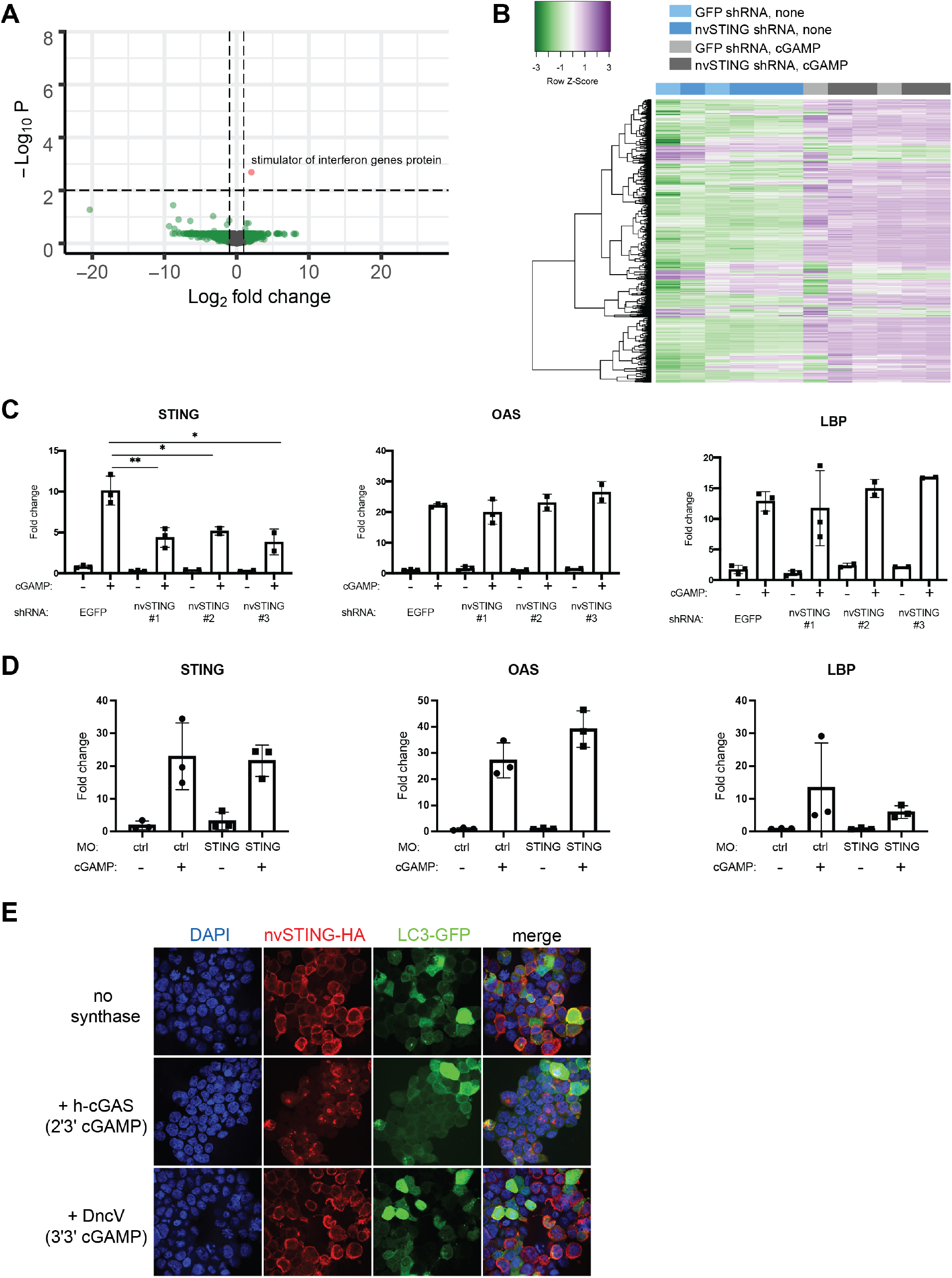
nvSTING knockdown does not impact the induction of genes by 2’3’-cGAMP. A) Volcano plot showing differential gene expression in 48 hour embryos treated with 2’3’-cGAMP that were injected with GFP shRNA or nvSTING shRNA. Positive fold-change indicates higher expression in GFP shRNA injected embryos. B) Clustered heatmap showing the expression of the top 1000 varied genes by RNA-Seq between embryos injected with either GFP or nvSTING shRNA and either untreated or treated with 2’3’-cGAMP. C) Fold change of nvSTING, nvOAS, and nvLBP assayed by Nanostring from experiments using 3 different shRNAs to knock down nvSTING expression. D) qRT-qPCR measuring genes of interest in 48-hour-old embryos injected with a control (ctrl) or nvSTING translation-inhibiting morpholino (MO) and treated with 2’3’-cGAMP. Fold changes were calculated as 2^-ΔΔCt^ and each point represents one biological replicate. Unpaired t test performed on ΔΔCt before log transformation; no significant differences. E) Immunofluorescence images of 293T cells transfected with plasmids encoding nvSTING-HA, LC3-GFP, and either empty vector, human cGAS or *V. cholera* DncV. Human cGAS is activated by the transfected DNA to produce 2’3’-cGAMP, and DncV, which produces 3’3’-cGAMP, is constitutively active in 293T cells.

**Figure S3:**
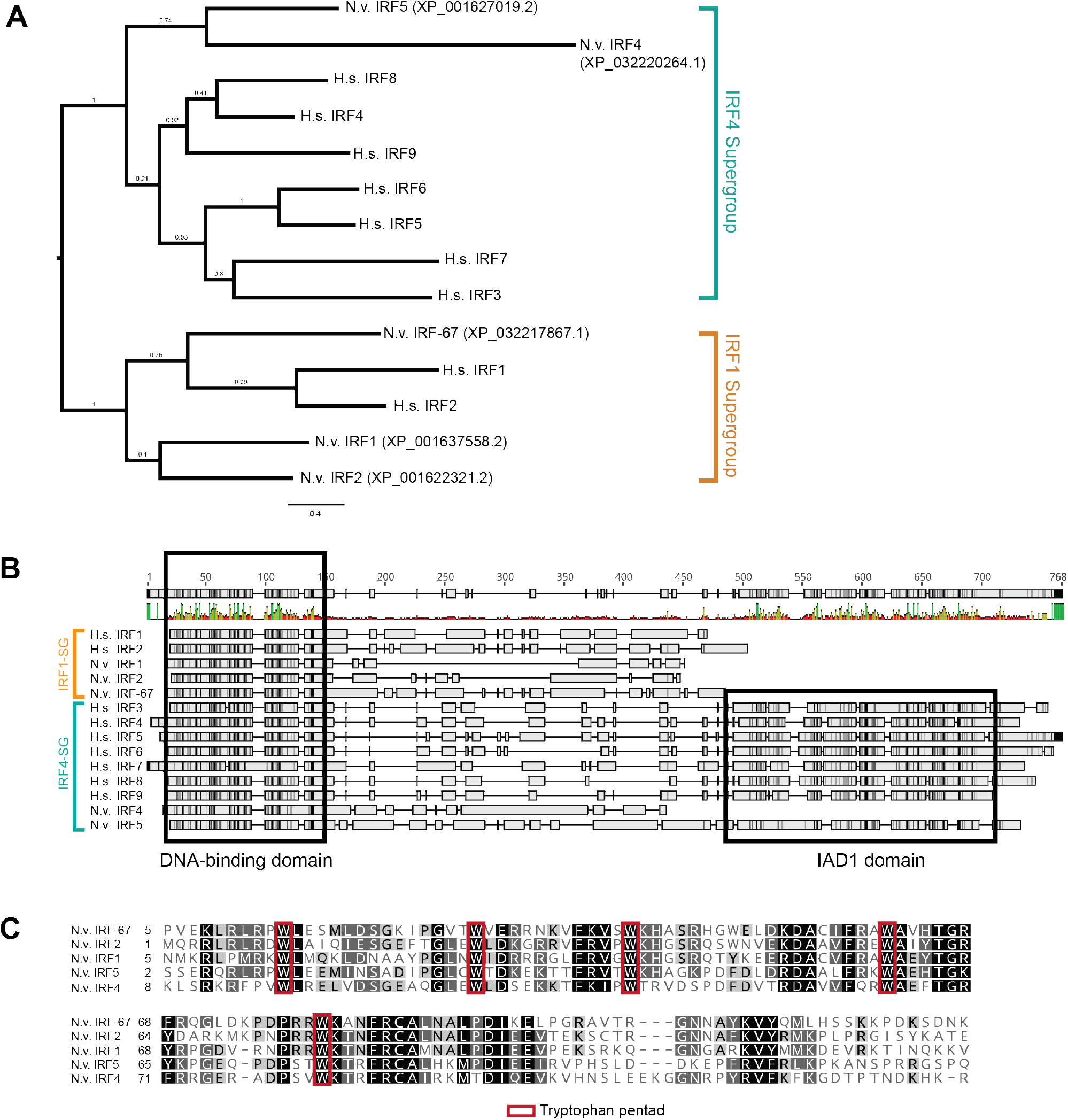
Phylogenetic study of *N. vectensis* IRFs. A) Phylogenetic tree of all human and *N. vectensis* IRF proteins. 3 nvIRFs cluster with members of the human IRF1 supergroup, while the other 2 nvIRFs cluster with the IRF4 supergroup. B) Full protein alignment of sequences in A). The DNA-binding domain is highly conserved between all *N. vectensis* and human IRFs. Among *N. vectensis* paralogs, only nvIRF5 contains an IAD1 domain. C) Alignment of all nvIRF DNA-binding domains with conserved tryptophan pentad outlined in red

**Figure S4:**
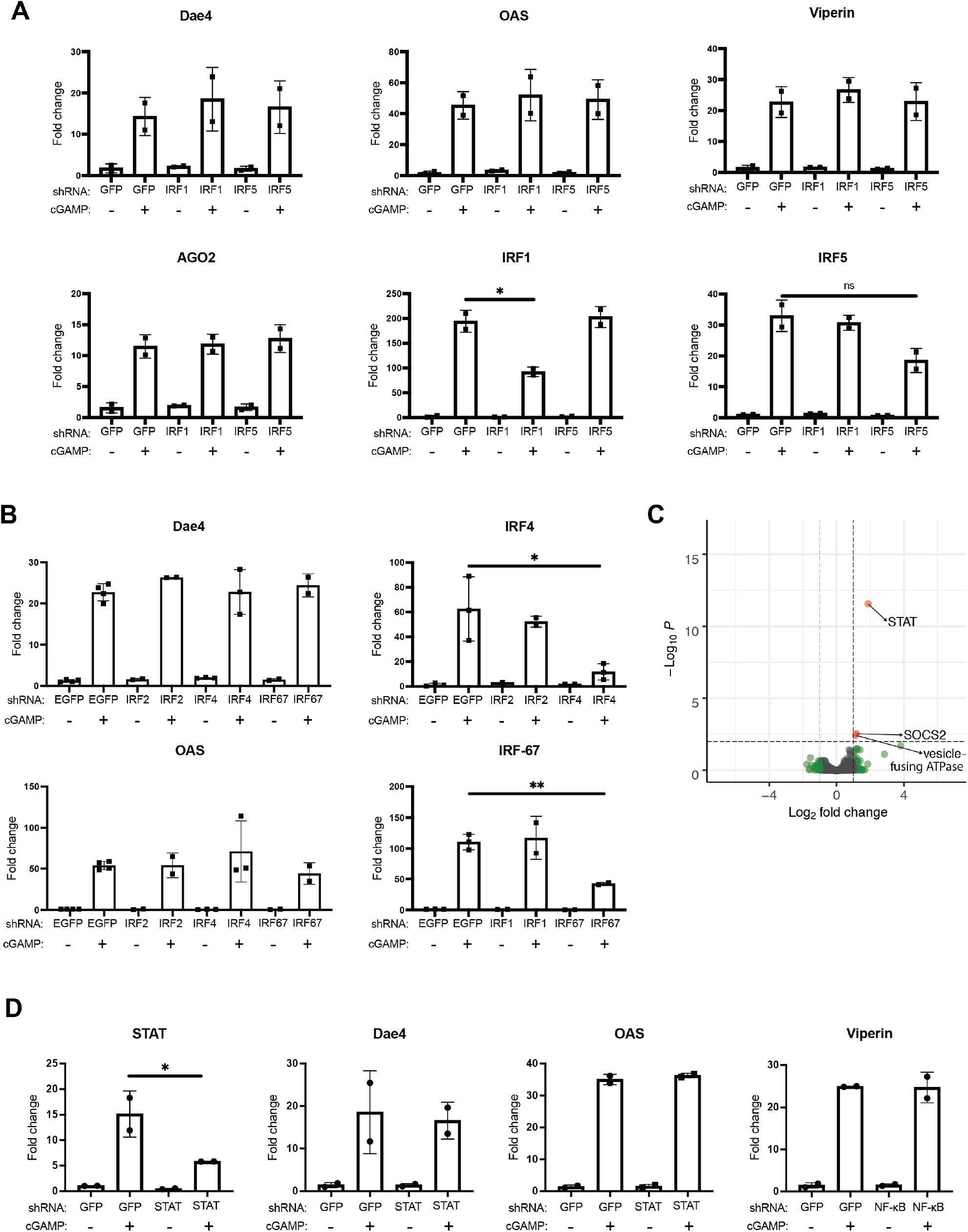
Knockdowns of nvIRFs or nvSTAT have no effect on 2’3’-cGAMP-induced gene expression. A) Fold changes in gene expression as determined by Nanostring in embryos microinjected with shRNAs targeting EGFP, nvIRF1, or nvIRF5 either untreated or treated with 2’3’-cGAMP. B) Fold changes in gene expression as determined by qRT-PCR in samples microinjected with shRNAs targeting EGFP, nvIRF2, nvIRF-67, or nvIRF4 either untreated or treated with 2’3’-cGAMP. Note that IRF2 is not induced by cGAMP and was mostly undetected in all samples; therefore it is possible that the knockdowns were unsuccessful. C) Volcano plot showing differential gene expression as determined by RNA-Seq in 48 hour embryos treated with 2’3’-cGAMP that were injected with GFP shRNA or nvSTAT shRNA. Positive fold-change indicates higher expression in GFP shRNA injected embryos. The GFP shRNA samples here are the same as those shown in Figure 2A. D) Fold changes in gene expression as determined by Nanostring in embryos microinjected with shRNAs targeting EGFP or nvSTAT either untreated or treated with 2’3’-cGAMP.

**Figure S5:**
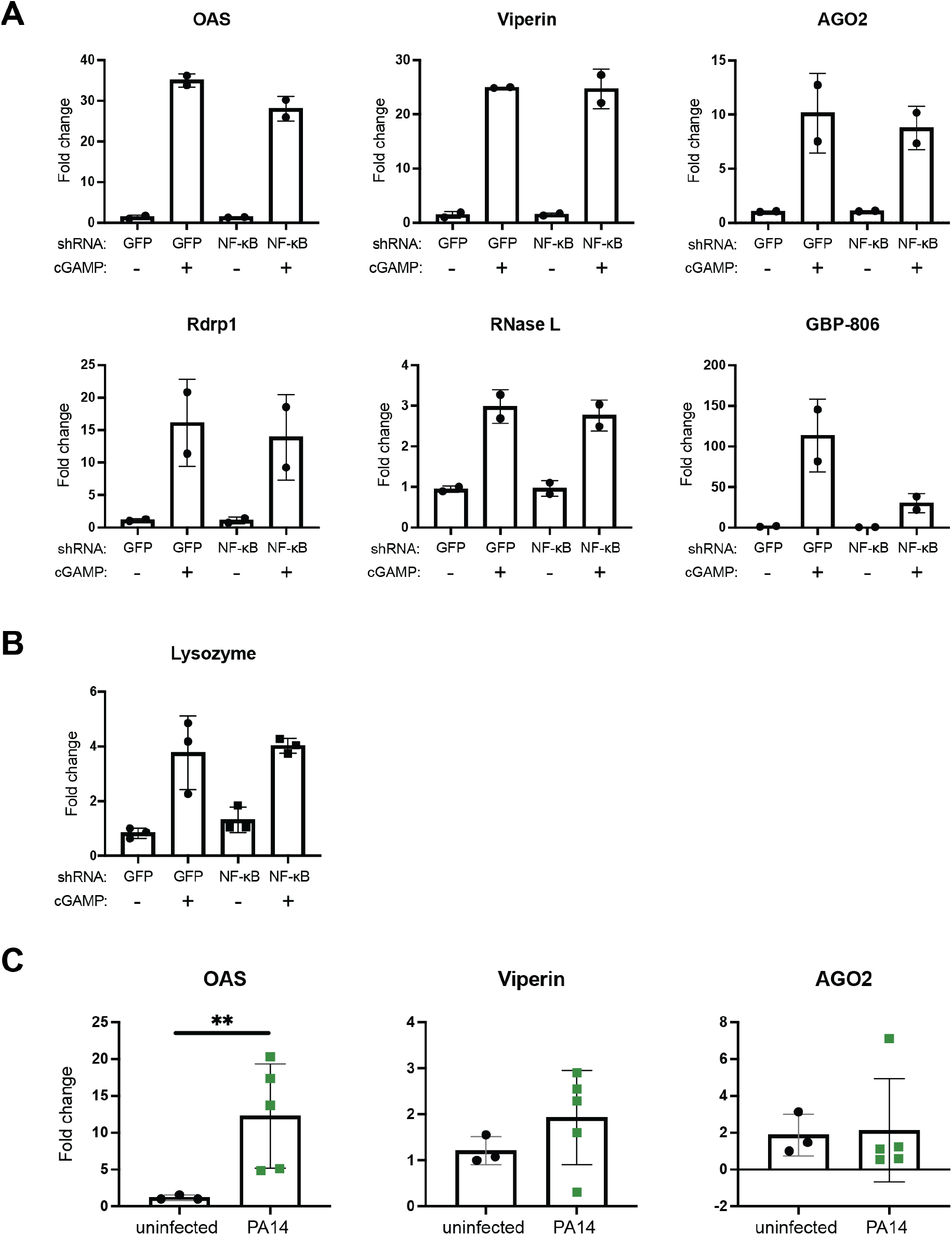
Anti-viral gene induction is not dependent on nvNF-κB. A) Fold changes in gene expression as determined by Nanostring in embryos microinjected with shRNAs targeting EGFP or nvNF-κB, either untreated or treated with 2’3’-cGAMP. The GFP shRNA samples here are the same as those shown in Figure S4. GBP-806 expression included to show anti-bacterial gene induction is lower in these samples. B) Fold changes in nvLysozyme expression as determined by qRT-PCR in embryos microinjected with shRNAs targeting EGFP or nvNF-κB, either untreated or treated with 2’3’-cGAMP. A+B) No significant differences in gene expression are observed between any cGAMP treated samples. C) qRT-PCR of putative anti-viral genes assayed at 48 hours post *Pa* infection (2 ×10^7^ CFU/ml). Each point represents one biological replicate; unpaired t test performed on ΔΔCt before log transformation. **p ≤ 0.01.

**Figure S6:**
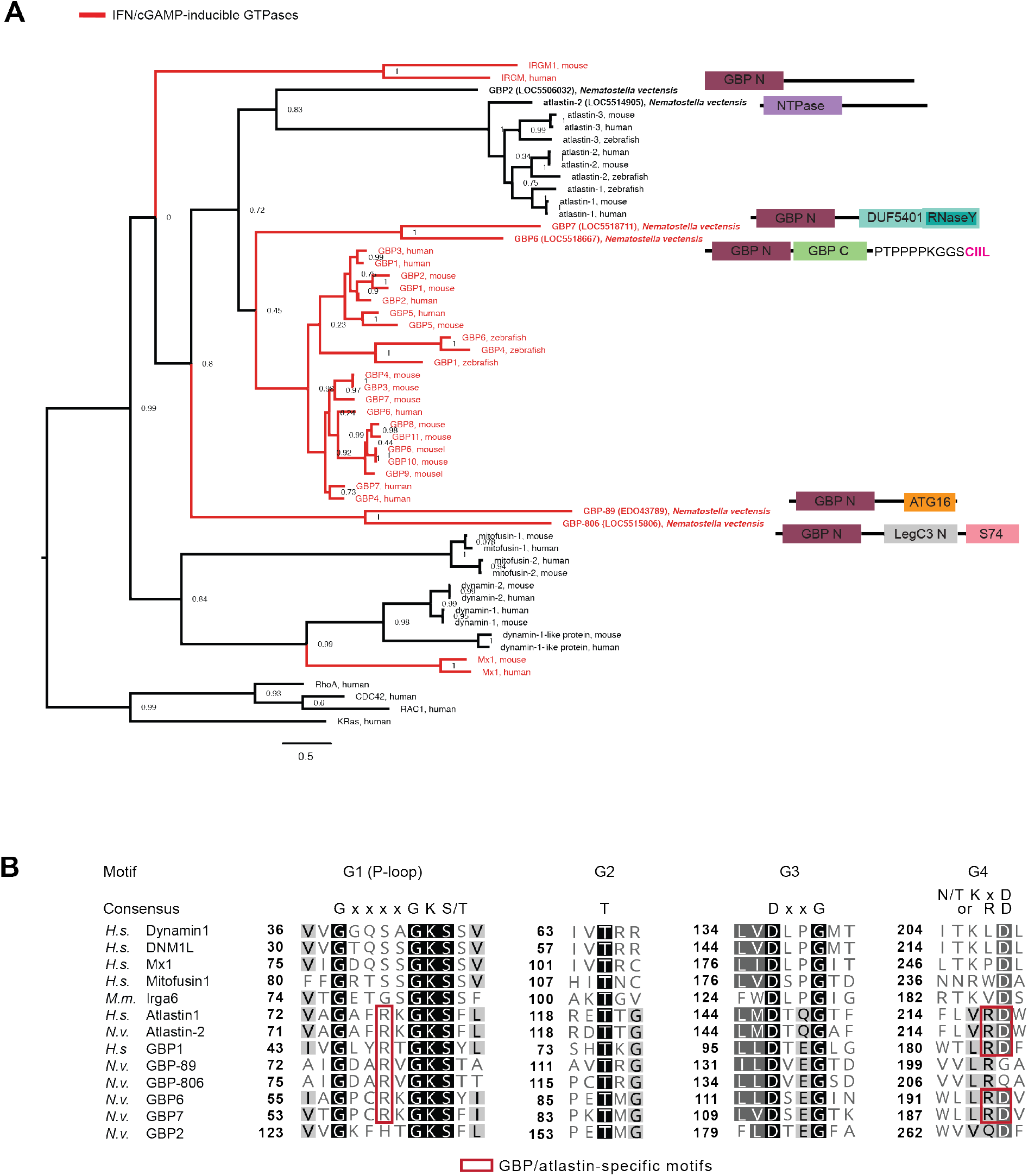
Phylogenetic study of *N. vectensis* GBPs. A) Phylogenetic tree of mammalian GTPases and putative *N. vectensis* GBPs made with the full protein sequences. Branches with mammalian interferon-induced GTPases and cGAMP-induced *N. vectensis* GBPs are colored red; these tend to cluster together. Domain structures of *N. vectensis* GBPs are displayed. B) Alignment of the GTPase domains of all *N. vectensis* GBPs and select mammalian GTPase. Conserved GBP and atlastin specific residues are highlighted in red.

**Figure S7:**
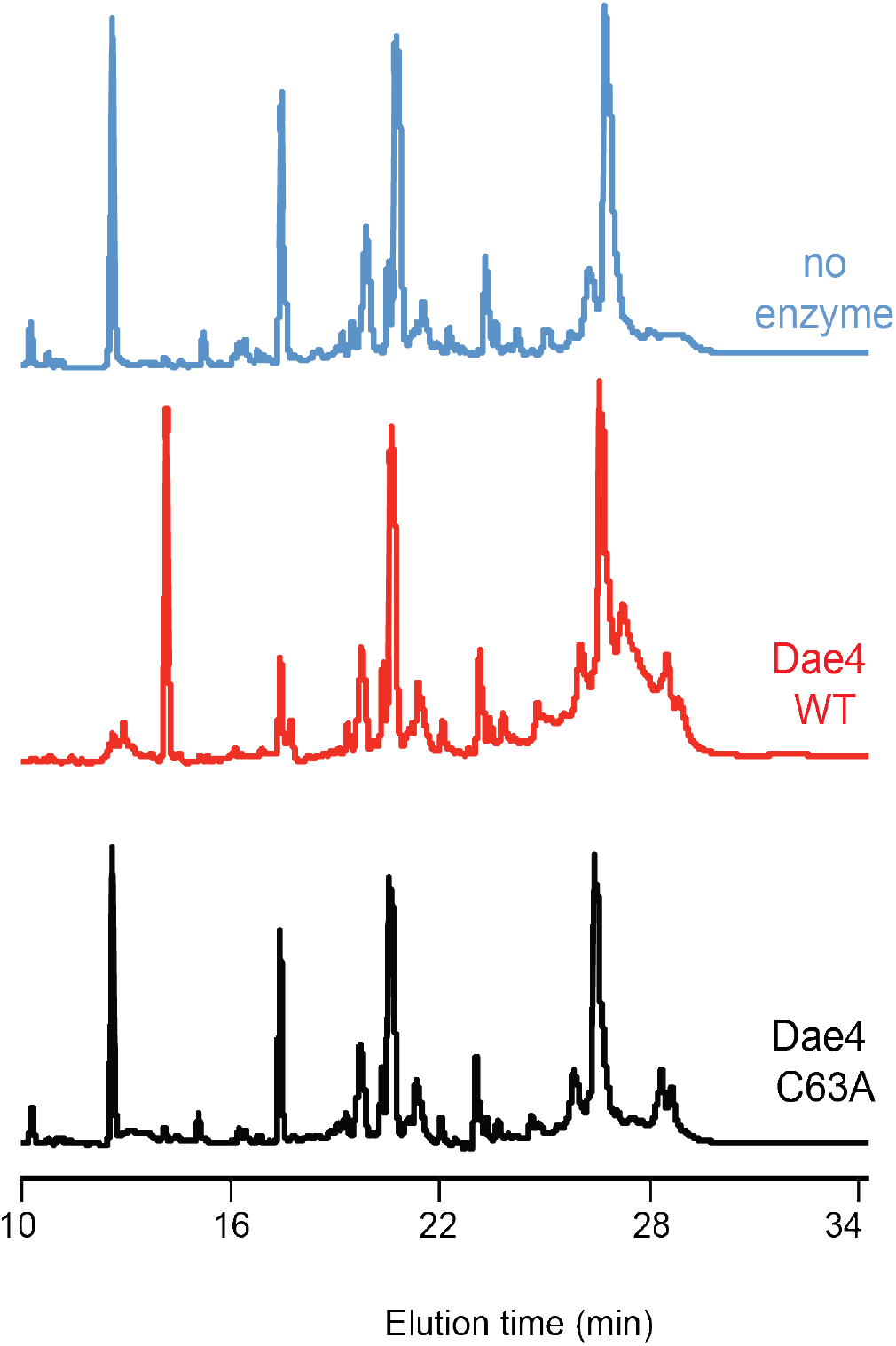
nvDae4 cleaves peptidoglycan from Gram-positive bacteria. Partial HPLC chromatograms of *Staphylococcus epidermis* peptidoglycan sacculi products resulting from incubation with buffer only (no enzyme), or 1 µM nvDae4 WT or C63A enzyme.

**Table S1:** Cnidarian-specific genes that are induced by 2’3’-cGAMP in an nvNF-κB-dependent manner.

| NCBI Gene ID | Domains | Only in Cnidaria? | Only in Anthozoa? | Homolog found in immune cells in *Stylophora pistillata* (1)? |
| --- | --- | --- | --- | --- |
| 5501851 | none | yes | yes | no |
| 5504224 | MDN1 super family | yes | yes | no |
| 5516219 | none | yes | yes | no |
| 5518710 | none | yes | yes | no |
| 116603205 | none | yes | yes | No homolog in *Stylophora pistillata* |
| 116603727 | none | yes | yes | No homolog in *Stylophora pistillata* |
| 116604070 | none | yes; except Bacillus spore coat proteins | yes; except Bacillus spore coat proteins | No homolog in *Stylophora pistillata* |
| 116604505 | none | yes | yes | No homolog in *Stylophora pistillata* |
| 116612667 | none | yes | yes | no |
| 116613998 | none | yes | yes | no |
| 116616875 | none | yes | yes; similar to 116620239 | No homolog in *Stylophora pistillata* |
| 116619128 | none | yes | no | yes |
| 116620239 | none | yes | yes; similar to 116616875 | No homolog in *Stylophora pistillata* |

**Dataset 1:** DESeq2 results for all RNA-Seq experiments

**Dataset 2:** All primers, oligos, and probes used in this study

